# B cell repertoire of children with atopic dermatitis exhibit altered IgE maturation associated with allergic food sensitization

**DOI:** 10.1101/2023.02.01.526538

**Authors:** Kirandeep K. Gill, Carolina Moore, Onyekachi Nwogu, John W. Kroner, Wan-Chi Chang, Jeffrey Burkle, Samuel J. Virolainen, Mariana L. Stevens, Asel Baatyrbek kyzy, Emily R. Miraldi, Jocelyn M. Biagini, Ashley L. Devonshire, Leah Kottyan, Justin T. Schwartz, Amal H. Assa’ad, Lisa J. Martin, Sandra Andorf, Gurjit K. Khurana Hershey, Krishna M. Roskin

## Abstract

IgE B cells produce antibodies responsible for the inappropriate specificity seen in allergic disease. In this study, we sequence antibody gene repertoires from a large, well-characterized early life cohort at risk for developing food allergy. We found that food allergen sensitization was associated with lower IgE mutation frequencies that was abrogated in children living with a pet dog, suggesting potential molecular mechanisms underlying the hygiene hypothesis (i.e., protective effects of pet ownership and non-antiseptic environs reported for allergic disease). We also observed increased IgE diversity and increased isotype-switching to IgE, suggesting that B cell development is altered in those with food allergen sensitizations relative to those without. IgE diversity was even more extreme in subjects who were highly likely to be peanut allergic. Unlike food antigen sensitization, we detected no effect of aeroallergen sensitization on IgE mutation levels and diversity. Consistent patterns of antibody rearrangement were associated with food allergen sensitization. These results have important implications for the allergic response driven by mast cells and basophils and the underlying drivers of the atopic march.

## INTRODUCTION

The antibodies secreted by IgE B cells drive allergy specificity and atopic disease^*1*^. We set out to study alterations in the B cell repertoire and its association with allergic sensitization in a cohort of children with atopic dermatitis (AD), a common chronic inflammatory skin condition and a major risk factor for allergic disease.

AD typically presents as dry skin and erythematous pruritic lesions. Nonlesional skin is also involved and likely promotes development and severity of atopic disease via subclinical barrier dysfunction and inflammation^*2,3*^. This skin barrier dysfunction contributes to the inappropriate immune response that develops in atopic diseases, including asthma and food allergy^*4*^. Mouse models suggest that immune dysregulation in the skin can also lead to inappropriate immune responses at other epithelial surfaces, such as the gut^*5*^. Taken together, these results suggest that barrier dysfunction is associated with wide-scale immune dysregulation. Out of this immune milieu, antibody producing B cells develop and mature.

When exposed to antigen or allergen, B cells clonally expand and activate somatic hypermutation (SHM) targeting B cell receptor (BCR) genes. While SHM correlates with greater antigen affinity of secreted antibodies, the role of SHM and affinity maturation in atopic responses is unclear^*6,7*^. Previous work characterized the BCR repertoires of children in Stanford’s Outcomes Research in Kids (STORK) birth cohort and found increased SHM levels in children with any form of atopy (asthma, hives, food allergy, and/or AD). However, the STORK study was limited in statistical power (only 17 out of 51 children had atopy)^*8*^. Thus, the generalizability of those findings is unclear, especially considering the wide spectrum of atopic disease severity. To further elucidate the immunological effects of allergic disease on the BCR repertoire, we performed high-throughput sequencing of rearranged immunoglobulin heavy-chain (IgH) genes from peripheral B cells of a subset of 147 children in the Mechanisms of Progression of AD to Asthma in Children (MPAACH) cohort^*9*^. MPAACH is the first US-based prospective early life cohort of children with AD. The cohort includes extensive participant phenotyping including several measures of AD severity and atopic disease progression, including measurement of allergic sensitizations.

Our data reveal changes in SHM levels and BCR diversity in children who have food allergen sensitizations as compared to children lacking food allergen sensitization. The change in diversity are more pronounced in subjects who were highly likely to be peanut allergic. By contrast, sensitization to seasonal or perennial aeroallergens had no statistically significant effect on the repertoire, suggesting that a different immune pathway leads to aeroallergen allergy development. Some of the allergen-associated BCR alternations were muted in children living with a pet dog, providing a possible molecular mechanism underling the protective effects of pet ownership reported for atopic disease and the hygiene hypothesis more broadly^*10*^. In children with food allergen sensitization, we also observed an increase in the overlap of the IgE compartment with other isotypes, suggesting that B cells are isotype switching to IgE at a faster rate in these subjects. Overall, our study associates major alterations in the maturation pathway of IgE B cells with food allergen sensitizations.

## RESULTS

### Antibody gene SHM increases with age across all isotypes, including IgE

As seen in other studies^*8*^, the early childhood antibody repertoire shows a progressive increase in SHM over time, which was statistically significant for all isotypes (Fig. 1A and fig. S1), all Bonferroni corrected *p*-values <5.4×10^-5^. Prior repertoire analyses either excluded IgE data or noted high variability in the IgE compartment preventing the detection of an age-associated increase in SHM^*8,11*^. Because of our large cohort (*n*=147, Table S1), we were able to detect an age-associated increase in SHM in the IgE compartment (Fig. 1A, Bonferroni corrected *p*-value of 5.4×10^-5^).

**Fig. 1:**
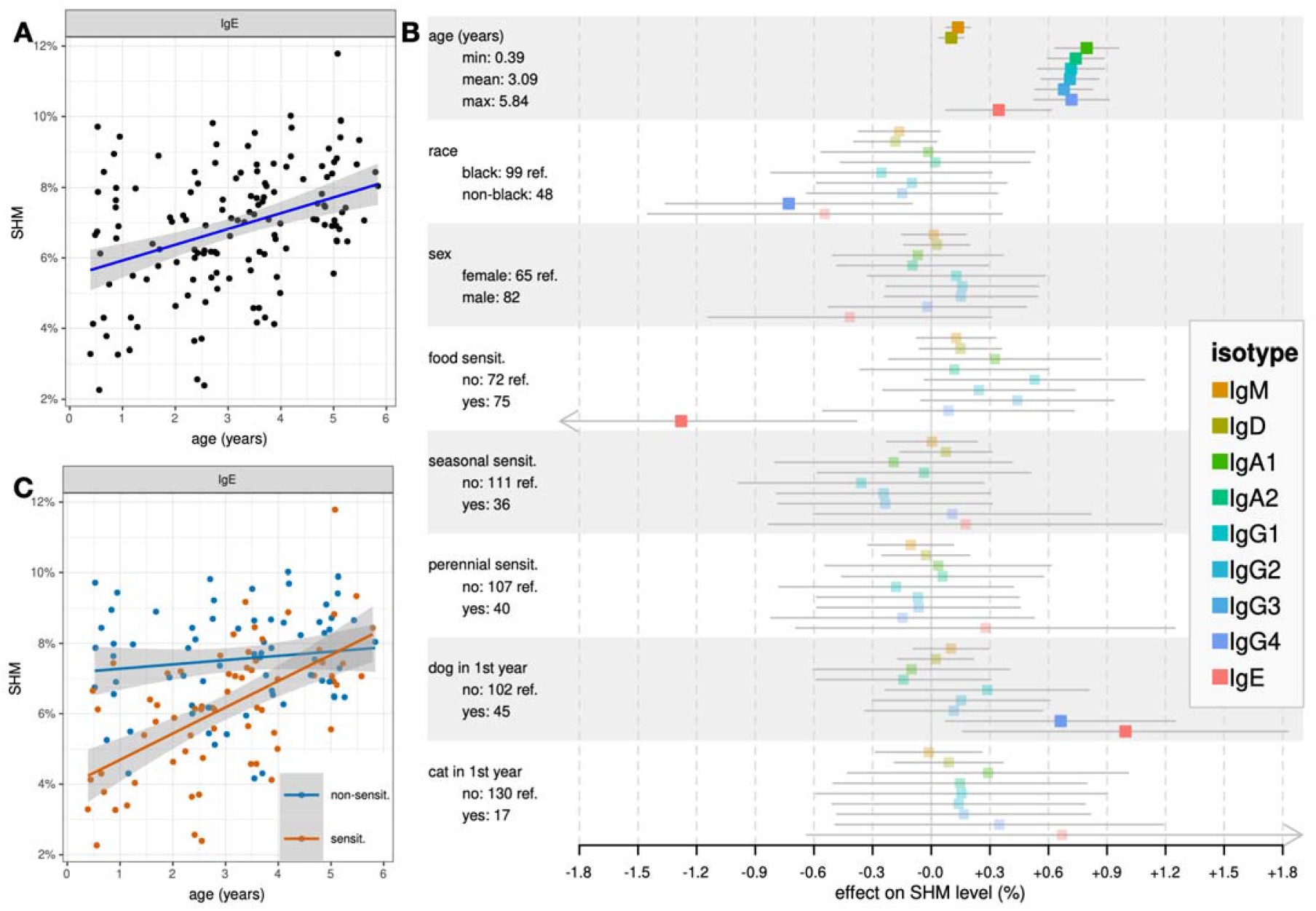
Linear modeling of somatic hypermutation (SHM). **(A)** An age-associated increase in SHM was seen for IgE and for the other isotypes (fig. S1). The shaded region represents the 95% confidence level from the linear model. **(B)** Multivariable linear regression modeling of SHM for all isotypes. Regression effect sizes (βs) for each predictor is shown on the *x*-axis in a forest plot/blobbogram. Confidence internals are adjusted for multiple hypothesis testing and statistically significant terms are highlighted. Age, in years, was modeled as a continuous variable; all other variables were categorical. The distribution of each variable is given in its row with the reference value of binary variables indicated. **(C)** Subjects with a food allergen sensitization had reduced IgE SHM across the study age range but this seems to resolve by age 6. Shaded regions represent the 95% confidence level from the linear models.

### Children with food allergen sensitizations had lower IgE SHM

To control for the age effect shown above, we include age as a covariant in a multivariable linear regression analysis of SHM levels with a Bonferroni correction (*m*=9) for the number of B cell isotypes tested^*12*^.

Sensitization to food allergens (positive SPT results to at least one of the thirteen food allergens tested, see Methods) was negatively correlated with IgE SHM (Fig. 1B). This relationship held across the age range of our cohort, with possible resolution in children ∼6 years of age (Fig. 1C).

### SHM not associated with measures of skin barrier function, AD severity, or variants in AD risk gene

Deep phenotyping was performed on a sub-cohort (*n=112*). Various measures of skin barrier function, AD severity^*9*^, and AD genetic risk^*13*^ were gathered. No significant differences in SHM were seen based on transepidermal water loss (TEWL), keratinocyte expression of filaggrin (FLG), presence of loss-of-function variants in FLG, or SCORing Atopic Dermatitis (SCORAD) measurements (fig. S2).

### A household dog is associated with higher IgE SHM, potentially countering the low IgE SHM associated with food allergen sensitization

The presence of a household pet, particularly dogs, has been proposed to attenuate the development of atopic disease in infants^*14,15*^. We found that living with a pet dog in the first year of life had a positive correlation with IgE SHM (Fig. 1B). Analysis of the effect on IgE SHM of the interaction between having a dog and food allergen sensitization shows that the decrease in IgE SHM associated with food allergen sensitization is limited to those *without* a household dog (reduction in IgE SHM of 1.89%, *p* < 5.2×10^-6^). Those living *with* a household dog in first year of life showed no food allergen sensitization related effect on IgE SHM (*p*=0.8).

No analogous effect was observed for cat ownership but with the small number of cat owners (*n*=17), we cannot draw strong conclusions.

### Repertoire diversity remains constant during early life while IgE and IgG4 diversity is greatly increased in food allergen sensitized subjects

Given the correlation between SHM and age (Fig. 1), we next examined if repertoire diversity also changes during early life. We measured repertoire diversity in such a way as to be orthogonal to SHM (see Methods). Unlike for SHM, we saw no correlation with age and repertoire diversity (Fig. 2, fig. S3A).

**Fig. 2:**
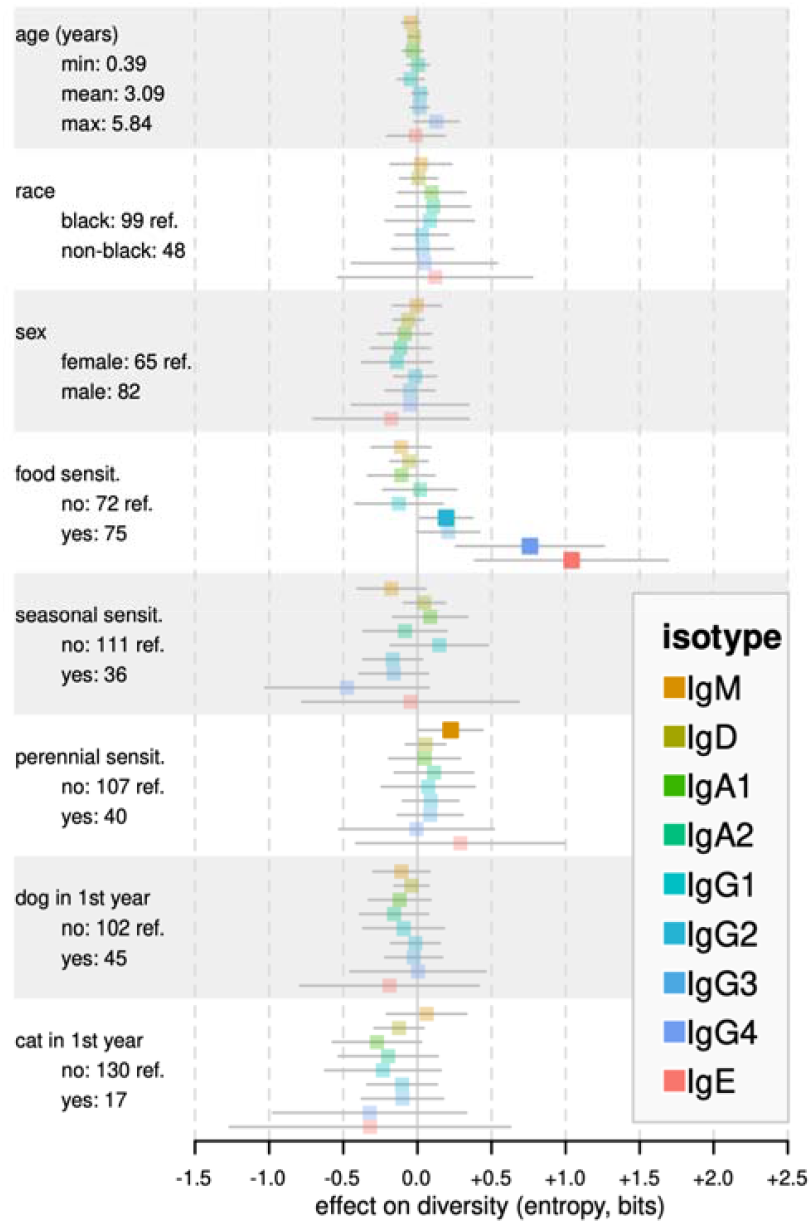
Linear modeling of repertoire diversity. Diversity, measured as the Shannon entropy of the of V- and J-segment usage frequencies, of the B cell clones expressing the given isotype. This measures combinatorial diversity and therefore is independent of the SHM differences described in Fig. 1. The regression effect size (βs) for each predictor is shown on the *x*-axis in a forest plot/blobbogram. Confidence internals are adjusted for multiple hypothesis testing and statistically significant terms are highlighted. Age, in years, is a continuous variable; all the others are categorical. The distribution of each variable is given in its row with the reference value of binary variables indicated.

Food allergen sensitized children relative to children without food allergen sensitization, had increased diversity in IgE, IgG2, and IgG4 (Fig. 2). IgE-positive B cells are relatively rare; thus, the baseline for IgE diversity is drastically lower than for the other isotypes. It is therefore striking that the median IgE diversity of food allergen sensitized subjects begins to approach the diversity observed in the other isotypes (fig. S3B).

### IgE B cell repertoire diversity correlates with serum IgE levels and is elevated in those highly likely to be peanut allergic

These changes in IgE diversity positively correlate with total serum IgE levels (*p*<4.2×10^-14^), suggesting that the peripheral IgE compartment is enriched for antibody secreting cells that are contributing to serum antibodies levels (Fig. 3A). Food allergen sensitization also had a significant effect on the association with serum IgE levels (p<0.0364, Fig. 3A). Subjects with peanut-specific IgE (sIgE) levels at or above 14 kU/L, which has a 95% positive-predictive value for peanut allergy^*16,17*^, had even higher IgE diversity (Fig. 3B). To control for age, IgE SHM levels were age-adjusted to the levels they would be expected to have at 3 years of age (the middle of the age range of this cohort) using a linear model (Fig. 1A). No differences in age-adjusted IgE SHM were observed in subjects highly likely to be peanut allergic (Fig. 3C). The analogous differences in IgE diversity and SHM in those highly likely to be milk allergic (fig. S4A-B)^16,18^ or egg white allergic (fig. S4C-D)^16,19^ were not statistically significate, though they trended higher for milk allergy.

**Fig. 3:**
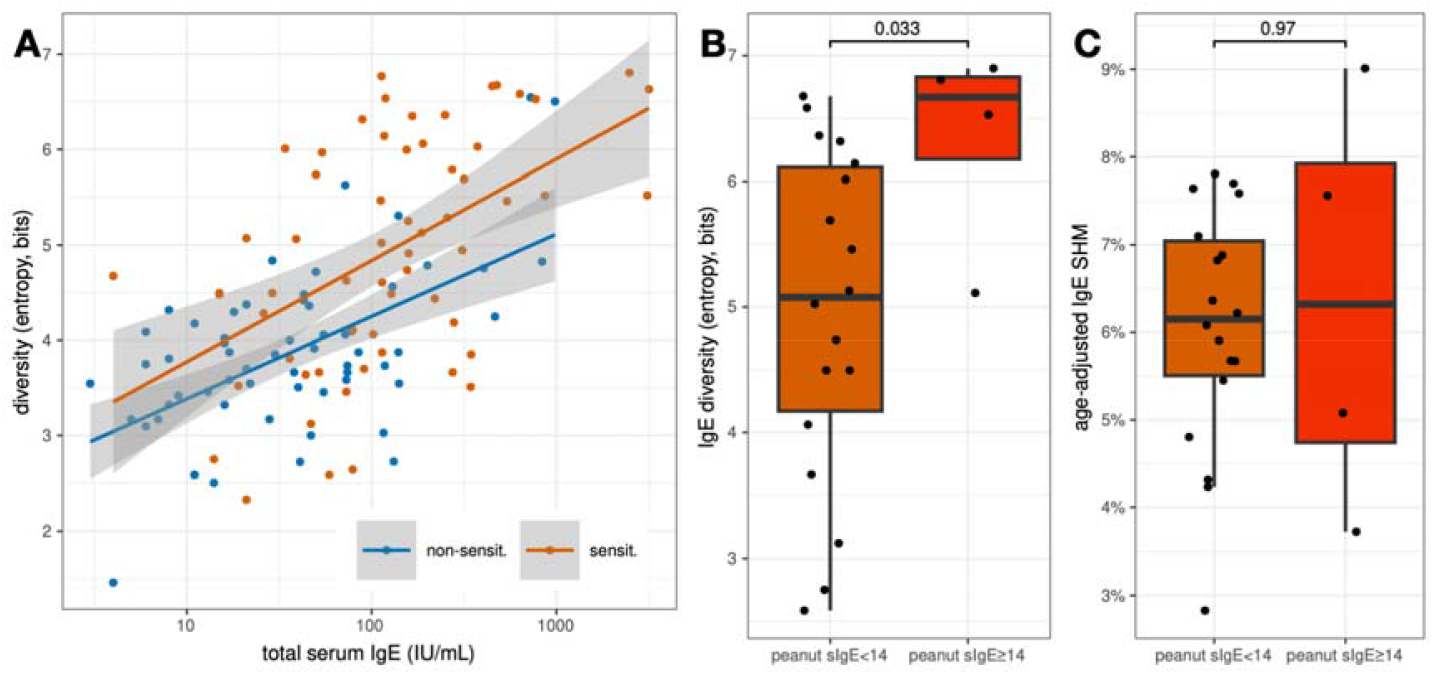
Diversity of the IgE□ B cell compartment correlates with serum IgE levels and is elevated in subjects highly likely to be peanut allergic. **(A)** Linear regression modeling showed a highly significant correlation of IgE□ B cell repertoire diversity with total serum IgE (log-scale, p < 4.2×10^-14^) with a food allergen sensitization having a significant effect (p<0.0364). **(B)** Subjects with peanut sIgE levels above 14 kU/L, estimated to be 95% predictive of peanut allergy by oral food challenge (OFC)^*16,17*^, exhibited even higher IgE diversity. **(C)** No change in age-adjusted IgE SHM was observed in those likely to be peanut allergic.

### IgE- and IgD-containing lineages have more overlap with class-switched isotypes in food allergen sensitized subjects relative to subjects without food allergen sensitizations

The increased diversity seen in the IgE compartment suggests an increase in isotype-switching to IgE. We looked for evidence of this by measuring the fraction of B cell lineages of a given isotype that contain clonal members of other isotypes. Compared to subjects with no food allergen sensitizations, food allergen sensitized subjects showed an increased frequency of overlap of IgE-positive clones with the other isotypes (Fig. 4). This increased overlap was also partially seen for IgD but was not seen in other isotypes, including IgM, the other “naïve” isotype.

**Fig. 4:**
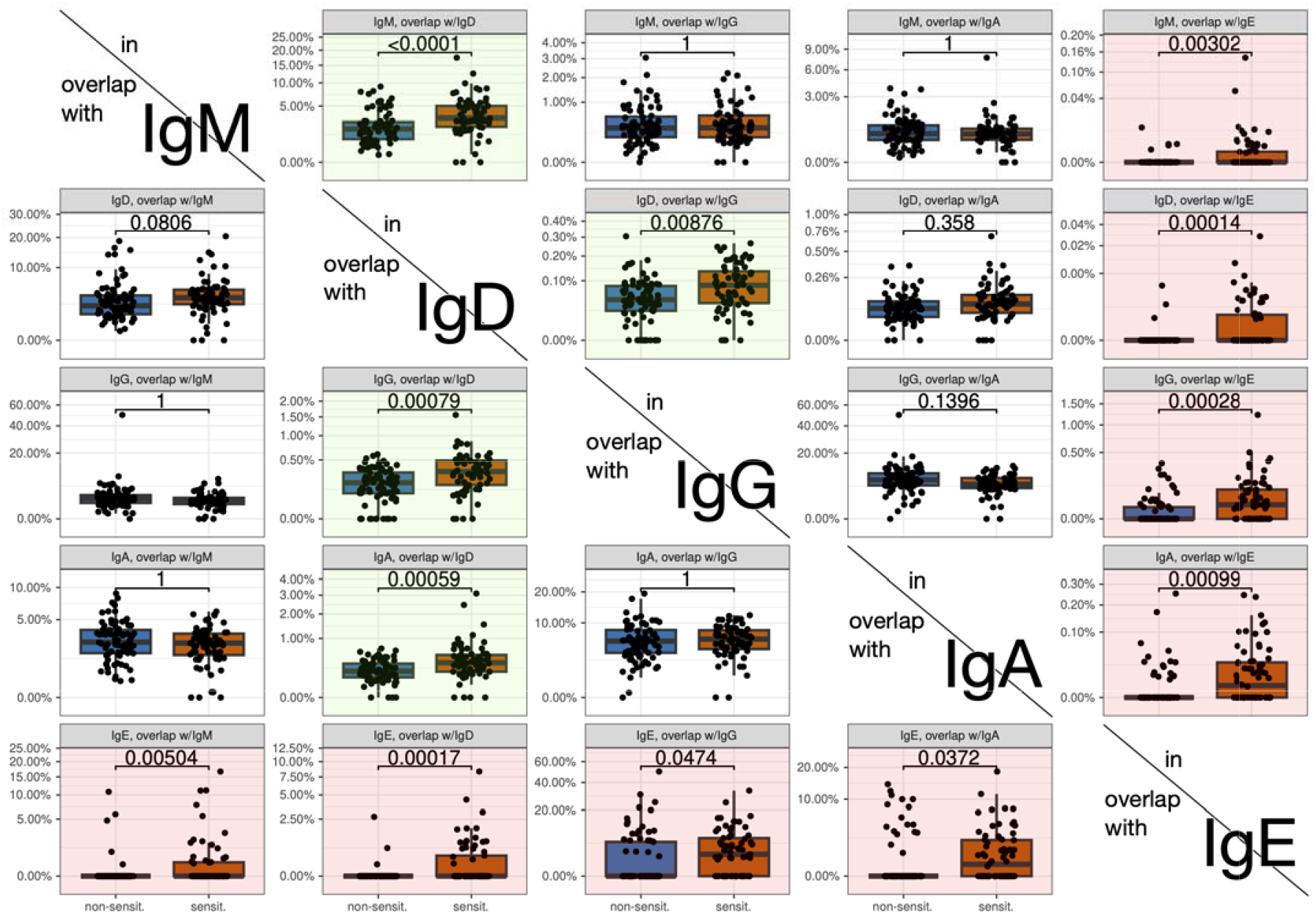
Clonal overlap between isotype compartments based on food allergen sensitizations status. For a given isotype, shown vertically, the per-subject fraction of clonal lineages of that isotype that also contain clonally related members of other isotypes, arrayed horizontally, is shown. The statistical significance for a difference in overlap based on food allergen sensitizations status was tested using a Wilcoxon test adjusted for multiple hypothesis testing. Statistically significance differences are highlighted in red for IgE and for green for IgD. All overlaps with IgE were statistically significantly increased in subjects with a food allergen sensitization as compared to those without such sensitizations. Several of the overlaps involving the IgD compartment showed a statistically significance increase in food allergen sensitized subjects. None of the overlaps involving non-IgE and non-IgD compartments were significantly different in food allergen sensitized subjects.

### Aeroallergen sensitization status is not detectably associated with changes in IgE SHM or diversty

Unlike for food allergen sensitization, we detected no changes in IgE SHM or diversity that were assoiated with aeroallergen sensitizations, either for seasonal or perennial aeroallergen (Fig. 1B and Fig. 2). With standard power calculation parameters (α=0.05, β=0.2) we are powered to detect changes in IgE SHM of 0.87% and diversity of 0.59 bits. These are smaller than the effect seen based on food allergen sensitization. Thus, we are powered to see an effect of aeroallergen sensitization that had a comparable effect size as seen for food allergens.

### Food allergen sensitized subjects exhibit skewed V-segment usage

Differences in SHM, diversity, and isotype overlap in subjects with food allergen sensitizations underlie complex, systemic changes in the immune state. Sequencing of the B cell receptor repertoire provides an opportunity to measure the effect of these changes on antibody repertoire. We examined the V-segment usage frequencies across each isotype of the subjects in our cohort. With 57 functional V-segments and 5 isotypes, this is a high-dimensional space^*20*^. We used principal component analysis (PCA) to linearly reduce this space to two dimensions capturing the largest trends in our data. Although PCA is unsupervised, i.e. does not take phenotype into account, the two axis of highest variance (PC1 and PC1) separate food allergen sensitized subjects from subjects with no food allergen sensitizations (fig. S5). We sharpened this discriminative power by using a L1-logistic regression model which was able to statistically separate food allergen sensitized subjects from subjects with no food allergen sensitizations (Fig. 5A). By looking at the regression weights (βs) we can identify the V-segments that are driving this separation. We see that, among others, usage of IGHV3-71 in IgG1 is strongly associated with not having a food allergen sensitization while use of IGHV3-69-1 in IgG2 is associated with being sensitized to food allergens (Fig. 5B).

**Fig. 5:**
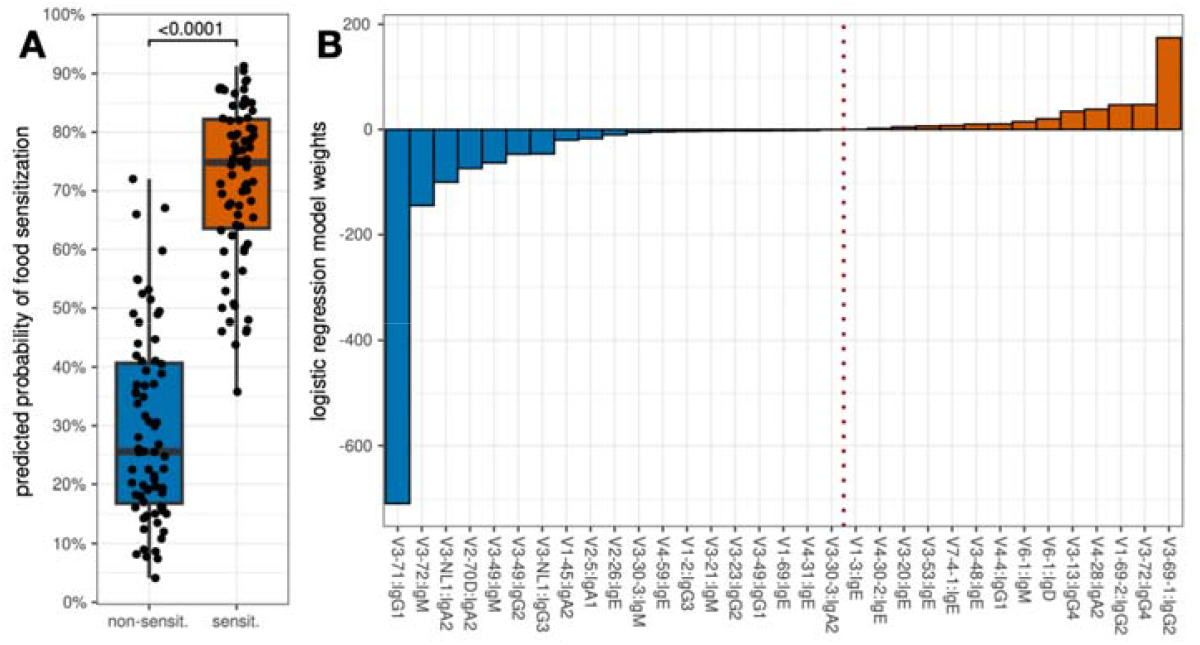
Logistic L1-regression model of V-segment usage. **(A)** We use logistic regression modeling of per-subject V-segment clonal usage frequencies in the 5 isotypes of each subject to predict food sensitization status. L1/LASSO regularization was used to fit a reduced model that showed excellent ability to statistically separate food allergen sensitized subjects from subjects with no food allergen sensitizations. **(B)** The non-zero feature weights from the reduced logistic regression model. Features with negative weight (blue) are associated with not having a food allergen sensitization while features with positive weight (red) are associated with being sensitized to food allergens. The dotted light separates the negative from positive feature weights.

## DISCUSSION

Our study provides insights into the dynamics of SHM during early childhood. Previous studies observed low levels of SHM during early life, extending as far back as gestational week 27^*11,21*^. The first major IgH repertoire study of infants (0-2 years) identified no increase in IgE SHM during early childhood^*22*^. This raised questions of how and when the developing immune system gains the additional 3-4% IgE SHM seen in healthy adults^*8*^. For the first time, using a large cohort, we identified age-dependent increases in IgE SHM from 0-6 years of age. As a corollary, immune receptor repertoire studies of children must control for age effects when considering differences in SHM. Controlling for such effects was critical to our analysis of atopic children.

Counter to our expectations, IgE SHM is decreased and potentially suppressed in children with food allergen sensitizations compared to unsensitized subjects. (Fig. 1B and Fig. 1C). A previous study reported *increased* SHM associated with atopic disease^*8*^, but this result presented a paradox for the *hygiene hypothesis*, which posits a protective role for increased antigen exposure^*23*^. If the “protective” antigen exposure results in increased SHM, why was increased SHM also associated with atopic disease? With a larger cohort, greater statistical power, and better control for confounders (e.g., age), our findings are consistent with the *hygiene hypothesis*: we find that food allergen sensitization is associated with a *decrease* in IgE SHM, i.e., an immune-repressed IgE milieu instead of a stimulated one. Our data also provide a possible explanation, at the molecular level, for the protective effects of dog ownership seen in epidemiologic studies, as pet ownership increases antigen load^*15*^. The presence of a household dog counters the reduction of IgE SHM levels associated with food allergen sensitization, thus helping maintain “normal” IgE SHM levels (Fig. 1B).

Protective effects of higher IgE SHM have also been reported in the context of specific immunotherapy (SIT) for allergic desensitization. Over the course of SIT, the IgE repertoire showed increased SHM, as did allergen-specific sequences, identified by phage display^*24*^. Thus, correction of low IgE SHM may be an important mechanism of SIT.

For food allergen sensitized subjects, the increased repertoire diversity in IgE (Fig. 2) together with that compartment’s lower SHM levels (Fig. 1B-C) suggests that B cells are isotype switching to IgE more readily in these subjects, before they can accumulate SHM. This is consistent with the increased percentage of IgE-positive clones that also contain members of other isotypes seen in food allergen sensitized subjects (Fig. 4). Early IgE switching is also supported by in-vitro functional studies, in which B cells from subjects with asthma or AD have a higher propensity to isotype-switch relative to B cells from non-atopic subjects^*25*^. Hoof et al. argued that allergen-specific IgG□ memory B cells rapidly switch to IgE and turn into antibody secreting cells upon allergen exposure^*26*^. The association in our data between food sensitization and the frequency of certain V-segments in IgG1 and IgG2 (Fig. 5B) is consistent with IgE serologic “memory” being stored in the IgG□ B cell compartment ^*26*^.The strong correlation in our data between IgE diversity and serum IgE antibody levels supports the idea that these newly switched cells are a major source of serum IgE antibodies (Fig. 3A).

We also observe increased overlap of IgE and IgD BCR sequences with other compartments, suggesting a shared pathway in IgD and IgE B cell development. A shared developmental pathway is also supported by correlation between aeroallergen-specific IgD and IgE antibodies^*27–29*^. An increase in extrafollicular IgE B cells in food allergen sensitized subjects could underlie the IgE SHM differences seen here. Dahlke and co-authors proposed an atypical developmental path for IgE□ B cell, based on analysis of IgE SHM^*30*^. They hypothesize that IgE B cells undergo reduced affinity selection relative to IgG B cells due to an extrafollicular origin of IgE B cells. It has been suggested that, in AD, CD27□IgE□ B cells are derived in a T cell-independent manner resulting in low SHM levels^*31*^. Further studies that link the B cell phenotype of these cells with their isotype and SHM status are needed. Such studies could also identify druggable vulnerabilities in this potentially pathogenic cell population, providing new therapeutic avenues for atopic diseases.

The influx of diverse B cells into the IgE compartment could be due to broad nonspecific, immune stimulation from a superantigen, such as those expressed by *Staphylococcus aureus* and implicated in AD^*32*^. While we do observe a broad systemic immune response, we also observe specific V-segment usage patterns that correlate with food allergen sensitization (Fig. 5B). Selection for these stereotypic rearrangements suggests that antigen selection is contributing to this response. Epidemiological studies find the protective effects of farming environments on atopic disease is restricted to specific livestock and farming activities, also suggesting some antigen-specific component^*33*^.

Several scales of AD severity have been proposed to measure clinical severity and skin barrier function. These clinical measures are not associated with changes in IgE SHM (fig. S2). Thus, the B cell receptor repertoire changes described here capture novel immunological correlates, that could be combined with other predictive features to identify endotypes of allergic disease and improve clinical outcomes.

The MPAACH cohort is an AD cohort. Further studies are needed to determine how these relationships hold for other cohorts and demographic groups. The repertoire changes described might be specific to this high-risk cohort. A larger study with non-AD subjects, particularly non-AD subjects with food allergen sensitizations, will provide valuable comparator groups.

The differences in IgE SHM described above, between food allergen sensitized subjects and subjects with no food allergen sensitizations, decreases with age and appears to dissipate around 6 years of age (Fig. 1C). This is consistent with natural history studies of food allergen sensitization and food allergy^*34–38*^. The repertoire data in this study are cross-sectional. Longitudinal data would elucidate the dynamics of this process and help determine if the immunophenotypes observed here are durable, or wax and wane with allergen sensitization.

The allergic response begins with degranulation and histamine release from mast cells and basophils. This requires IgE antibodies against multiple epitopes to bring IgE-receptor complexes on these cells into proximity to trigger crosslinking and downstream signaling^*39,40*^. Christensen and colleagues studied the combinatorial effect of dust mite allergen-specific IgE antibodies on basophil degranulation^*41*^. They found that a mix of high and low affinity antibodies were as potent as two high affinity antibodies and that the largest effect was seen with a diverse mix of antibodies. This suggests an interplay between IgE SHM, and the associated increase in antibody affinity, and IgE antibody diversity.

The increases in IgE repertoire diversity associated with food allergen sensitization was even more extreme in subjects who were highly likely to be peanut allergic based on their serum peanut-specific IgE levels (Fig. 3B). This suggests that peanut allergy is a similar but more extreme form of peanut allergen sensitization, as opposed to a categorically different immune response. No change in IgE SHM was observed based on a high likelihood of peanut allergy (Fig. 3C). This argues that diversity in the IgE compartment is much more important than IgE affinity (as measured by SHM) when it comes to the peanut allergy. How these findings generalize to other foods remains to be seen as the numbers of subjects with egg or milk sensitization was low. Additionally, showing how these findings translate to people with proven food allergy, by oral food challenge, will be the subject of future studies.

In summary, our analysis of the BCR repertoire in a cohort of AD subjects finds large immune perturbations associated with food allergen sensitization. We observe lower IgE SHM in subjects with food allergen sensitizations compared to those without food allergen sensitizations. This result provides explanation for the protection against atopic diseases conferred by pet ownership and non-antiseptic living that has been described epidemiologically. In food allergen sensitized subjects, we also see increased diversity in the IgE compartment, and this increase was even higher in those likely to be peanut allergic. We also see increase overlap between the IgE compartment and other isotypes. This suggests that B cells are isotype-switching to IgE at a faster rate in these subjects and the resulting increase in diversity is playing an important role in food sensitization and allergy. These results quantify the immune perturbations in subjects with food allergen sensitization and could be deployed in the future to aid in diagnosis and risk stratification of allergic disease.

## HMATERIALS AND METHODS

### Study design

The Mechanisms of Progression of AD to Asthma in Children (MPAACH) is a prospective early life cohort of children with AD who are followed with yearly visits^*2*^. Inclusion criteria were (1) age ≤2 years at enrollment, (2) gestation of at least 36 weeks and (3) a diagnosis of AD (based on the Hanifin and Rajka criteria for AD^*42*^) or the parent(s)/legal authorized representative indicates a positive response to each of the 3 questions from the Children’s Eczema Questionnaire^*43*^. Exclusions criteria include (1) a comorbid lung condition including cystic fibrosis, congenital anomaly, or bronchopulmonary dysplasia, (2) dependence on immunosuppression or oral steroids for a medical condition other than asthma, (3) condition that precludes sampling of the proposed biologic samples or completion of spirometry, and (4) a bleeding diathesis. This study was approved by the Institutional Review Board at Cincinnati Children's Hospital Medical Center (CCHMC). All subjects or their legal representative signed informed consent documents prior to participation. The study population in this analysis is a subset of size *n*=147 of MPAACH enriched for those sensitized to peanut, hen egg, or cow milk. The characteristics of this sub-cohort are shown in Table S1.

### Atopic dermatitis phenotyping

AD severity was evaluated clinically using SCORing Atopic Dermatitis (SCORAD)^*44*^. Skin barrier integrity was quantified using transepidermal water loss (TEWL) as measured on lesional skin using the DermaLab TEWL probe (Cortex Technology, Hadsund, Denmark). Skin barrier dysfunction was quantified by expression of filaggrin (FLG) from lesional keratinocytes normalized to the expression of the 18S rRNA, as described previously^*2,45*^. Serum IgE levels were quantified via ImmunoCap on a Phadia 250 Immunoassay Analyzer (ThermoFisher Scientific, Waltham, Wash). Germline sequencing and variant calling of the FLG gene was performed using targeted deep sequencing and the Genome Analysis Toolkit (GATK) HaplotypeCaller^*46*^ version 4.2.0.0 following best practices as described in ref. *(13)*.

### Allergic sensitization

Allergic sensitization was measured on the day of blood draw by skin prick testing (SPT) with a panel of 13 foods (milk, peanut, egg white, egg yolk, cashew, almond, walnut, pistachio, pecan, hazelnut, Brazil nut, wheat, soy) foods and 11 aeroallergens that included 7 seasonal (two tree mixes, ragweed, two mold mixes, grass, weeds) and 4 perennial (dog, cat, cockroach, mites) allergens^*2*^. A positive SPT result was a wheal diameter >3 mm larger than the diluent control.

### IgH sequencing libraries

Peripheral blood mononuclear cells (PBMCs) were isolated from whole blood using the SepMate column system (StemCell Technologies) with Histopaque-1077. AllPrep column purification (Qiagen, Valencia, CA) was used to isolate RNA. Complementary DNA (cDNA) was generated from total RNA using SuperScript III (Invitrogen, Waltham, MA) and random hexamers. PCR amplification of IGH rearrangements from the cDNA template was carried out according to ref. ^*8*^. In brief, template was amplified using multiplexed primers targeting IGHV gene segments using the BIOMED-2 first framework primers and isotype-specific primers located in the CH1^*47,48*^. This first round PCR primers also included half of the Illumina P5 and P7 adapter sequences. The first round PCR used Platinum PCR Master Mix (Applied Biosystems, Waltham, MA) according manufacture instructions, and the following program: 95°C for 2 min, 35 cycles of (95°C for 30 sec, 60°C for 90 sec, 72°C for 60 sec), and a final extension at 72°C for 10 min. Illumina adapters were completed by a second PCR carried out with the Qiagen Multiplex PCR kit (Qiagen, Valencia, CA), using 1 μl of the first PCR product as the template in a 20 μl reaction with the following program: 94°C for 15 min, 12 cycles of (94°C for 30 sec, 60°C for 45 sec, 72°C for 90 sec), and a final extension at 72°C for 10 min. Each isotype was amplified separately to decrease chimeric product generation. PCR reactions for all samples were pooled and purified by agarose gel electrophoresis and gel extracted using the QIAquick kit (Qiagen, Valencia, CA). Sequencing of the final libraries was performed on the Illumina MiSeq instrument using 600-cycle kits by the CCHMC DNA Core.

### IgH sequence annotation

High-throughput sequencing data was processed as previously described^*8,49*^. In short, 300bp paired-end reads were merged using FLASH^*50*^. Reads were mapped to samples using barcode sequences. The V-, D-, and J-segments and framework and CDR regions were identified using IgBLAST (version 1.16.0)^*51*^. The germline sequences of the V-, D-, and J-segments were taken from the international ImMunoGeneTics information (IMGT) database (Aug. 2022 version)^*20*^. Sequences were quality filtered to include only productive reads with a CDR-H3 region, minimum V-segment alignment score of 70, and minimum J-segment alignment score of 26. The isotype of each transcript was determined by exact matching to a database of constant region gene sequences upstream from the primer^*20*^. Subjects were required to have at least 5 clones of a given isotypes or were excluded from the analysis due to poor estimates of per-subject mean SHM, diversity, or V-segment usage.

### Clonal inference

Clonal relationships were assigned as previously described, using mmseqs2 (Sep. 2019 build)^*52*^ to cluster the sequences into clones using the same V- and J-segments (without considering the allele), equal CDR-H3 length, and at least 90% CDR-H3 nucleotide identity^*48*^. Mutation levels and V- and J-usage frequencies are calculated per clone using this inference of clonal grouping.

### SHM calculation

The per-read SHM level was calculated as the percentage of V-segment bases that did not match the inferred germline sequence, excluding the region targeted by primer sequences and the part of CDR3 encoded by the end of the V-segment. For each isotype, the clonal SHM level was calculated as the median SHM level of all reads of that isotype in clonal lineage. For each isotype, the subject SHM level was calculated as the mean SHM of each lineage of that isotype.

### Diversity analysis

For each isotype and subject, the isotype-specific usage frequency of V- and J-segment pairs of each clone was calculated. From this distribution, Shannon entropy was calculated, in bits, to measure the diversity^*53*^. Since only extreme levels of SHM would cause IgBLAST to misclassify a V- or J-segment artificially increasing diversity, this measure is independent of SHM.

### Statistical analysis

Statistical analysis and graphs were generated using the R statistical language (version 4.4.0), in RStudio (build 999)^*54,55*^. Linear regression modeling was performed using the lm R function. All confidence intervals from the linear models were adjusted with a Bonferroni correction for multiple hypothesis testing^*12*^. The statistical significance of population differences was tested using the two-side Wilcoxon test as implemented by the wilcox.test function in R and was also corrected for multiple hypothesis testing with a Bonferroni correction^*12*^. A p-value of 0.05 was considered significant. Plots were generated using the ggplot2 (version 3.5.1), ggpubr (version 0.6.0), forestplot packages (3.1.6)^*56–58*^. Box-whisker plots show median (horizontal line), interquartile range (box), and 1.5 times the interquartile range (whiskers).

Confidence intervals around regression curves show 95% confidence intervals and were calculated using the stat_smooth ggplot2 function. Confidence intervals around scatter plots were calculated using stat_ellipse with a level of 0.68 and using a multivariate t-distribution. Principle Component Analysis (PCA) was done using the pca function of the PCAtools R package (version 2.18.0)^*59*^. Logistic L1/LASSO regression was performed using the glmnet package (version 4.1-8)^60,61^. Regularization strength (λ) was selected by taking the value which minimizes the mean cross-validated error from 100 runs of cv.glmnet. For the PCA and logistic regression analysis, V-segment per isotype clonal usage frequencies that were rare (detected in fewer than 10% of subjects) were drop from the analysis.

## Acknowledgments

The authors thank all the children and their families who participated in the MPAACH cohort, the Schubert Research Clinic of Cincinnati Children’s Hospital Medical Center (CCHMC) for assistance with research participants, the CCHMC DNA Sequencing and Genotyping Core (in particular Brian Quinn for help optimizing library construction), the CCHMC Information Services for Research (IS4R) group for hosting the data storage and processing infrastructure.

## List of Supplementary Materials

Fig. S1. Linear modeling of somatic hypermutation (SHM).

Fig. S2. Linear modeling of SHM, skin barrier function, AD severity and genetic risk.

Fig. S3. Repertoire diversity across all isotype compartments.

Fig. S4: IgE diversity and age-adjusted IgE SHM is not significantly altered in subjects highly likely to be milk or egg white allergic.

Fig. S5. Component analysis (PCA) of V-segment usage.

Table S1. Demographic, clinical, and IgH sequencing characteristics of the cohort.

## Funding

National Institutes of Health (NIH) National Institute of Allergy and Infectious Diseases (NIAID) grant U19 AI070235 (GKKH, JBM, LJM, WCC), National Institute of Arthritis and Musculoskeletal and Skin Diseases (NIAMS) grant P30 AR070549 (LK, SA)

NIAID Asthma and Allergic Diseases Cooperative Research Centers (AADCRC) Opportunity Fund (KMR, SA).

Cincinnati Children’s Hospital Medical Center, Center for Pediatric Genomics (CpG) Pilot Grant (KMR).

Food Allergy Research and Education (FARE) Biobank and Biomarker Discovery Center (BBDC) (AHA, SA).

## Author contributions

Conceptualization: KG, SA, GKKH, KMR

Methodology: KG, CM, ON, SJV, MLS, ERM, JMB, LJM, SA, KMR

Investigation: KG, CM, ON, SJV, JWK, WCC, ABK, JTS, AHA, KMR

Visualization: KG, KMR

Funding acquisition: SA, GKKH, KMR

Project administration: JWK, WCC, MLS, JMB, KMR

Supervision: ALD, LK, JTS, AHA, GKKH, LJM, KMR

Writing, original draft: KG, SA, GKKH, KMR

Writing, review & editing: KG, CM, ON, JWK, WCC, SJV, MLS, ABK, ERM, JMB, ALD, LK, JTS, AMA, LJM, SA, GKKH, KMR

## Data and materials availability

All data associated with this study are available from the NIH Short Read Archive (SRA) under BioProject PRJNA857098. All python and R code is available from https://github.com/roskinlab/irad/.

