## Supplemental Materials for "B cell repertoire of children with atopic dermatitis exhibit altered IgE maturation associated with allergic food sensitization"

**Table S1.** Demographic, clinical, IgH sequencing statistics of the subjects in this cohort. *Numbers of rearranged IgH sequence reads and number of B cell clones (see Methods), also shown by isotype.

|  |  | food sensitization status | | |
| --- | --- | --- | --- | --- |
|  |  | non-sensit. | sensit. | all |
| n |  | 72 | 75 | 147 |
| age (years) | minimum | 0.53 | 0.39 | 0.39 |
|  | mean | 3.22 | 2.96 | 3.09 |
|  | maximum | 5.84 | 5.79 | 5.84 |
| sex | female | 37 | 28 | 65 |
|  | male | 35 | 47 | 82 |
| race | black | 56 | 43 | 99 |
|  | non-black | 16 | 32 | 48 |
| filaggrin expression (lesional) | mean | 9.52×10^-3^ | 5.24×10^-3^ | 7.39×10^-3^ |
|  | missing | 7 | 11 | 18 |
| filaggrin loss of function | yes | 4 | 13 | 17 |
| variants | no | 63 | 55 | 118 |
|  | missing | 5 | 7 | 12 |
| TEWL (lesional) | mean | 17.8 | 19.9 | 18.9 |
|  | missing | 3 | 2 | 5 |
| SCORAD | mean | 20.8 | 27.5 | 24.3 |
|  | missing | 1 | 0 | 1 |
| household pet | dog | 21 | 24 | 45 |
| (first year of life) | cat | 10 | 7 | 17 |
|  | both | 3 | 3 | 6 |
| IgH reads* | mean | 133,879 | 302,067 | 219,244 |
| IgM | mean | 49,098 | 80,866 | 65,166 |
| IgD | mean | 34,939 | 81,873 | 58,761 |
| IgA | mean | 25,816 | 52,027 | 39,119 |
| IgG | mean | 21,191 | 54,343 | 38,018 |
| IgE | mean | 3,463 | 34,996 | 19,369 |
| IgH clones* | mean | 60,446 | 122,982 | 92,187 |
| IgM | mean | 29,289 | 52,349 | 40,952 |
| IgD | mean | 21,331 | 51,031 | 36,405 |
| IgA | mean | 6,177 | 11,605 | 8,932 |
| IgG | mean | 5,575 | 12,764 | 9,334 |
| IgE | mean | 46 | 286 | 168 |

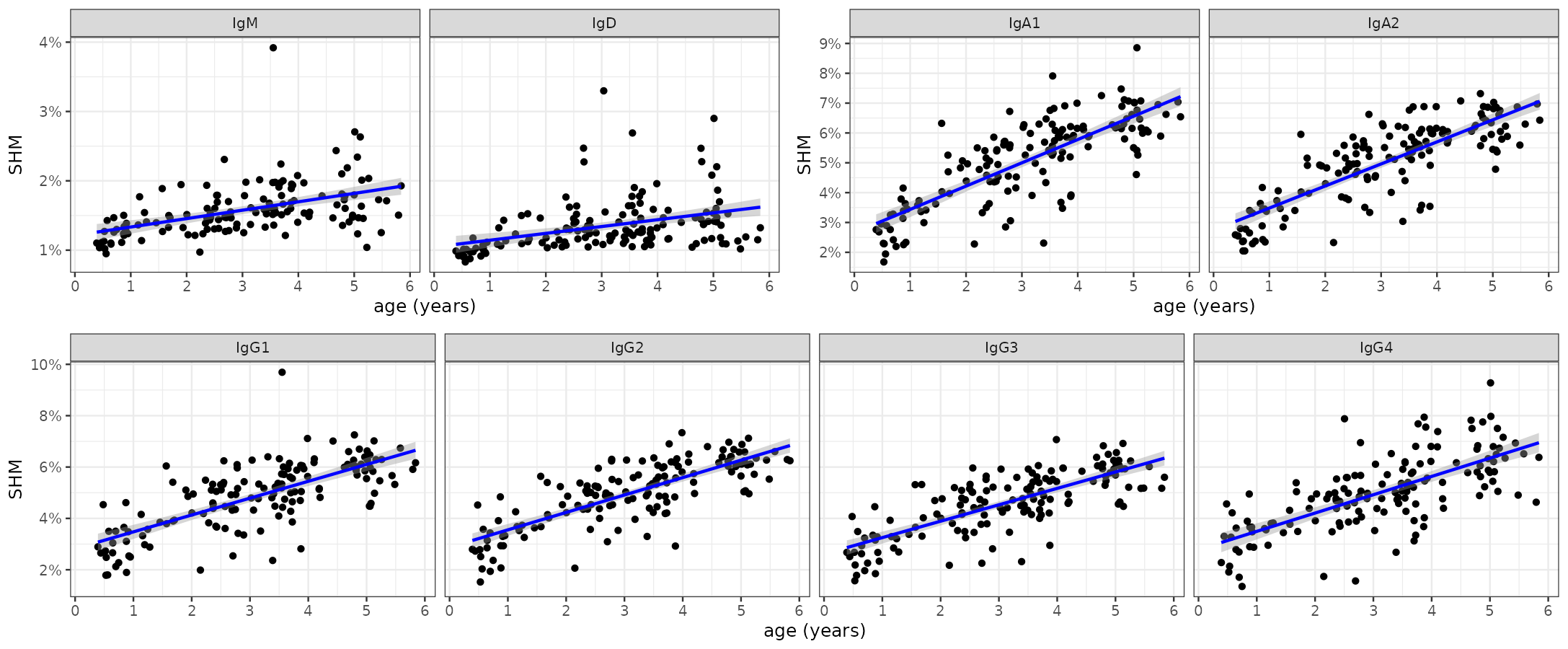

**Fig. S1: Linear modeling of somatic hypermutation (SHM).** An age-associated increase in SHM was seen for all isotypes. The shaded region represents the 95% confidence level from the linear model. Panel for IgE is shown in Fig. 1A. All models were statistically significant, Bonferroni corrected p-values < 5.4×10^-5^.

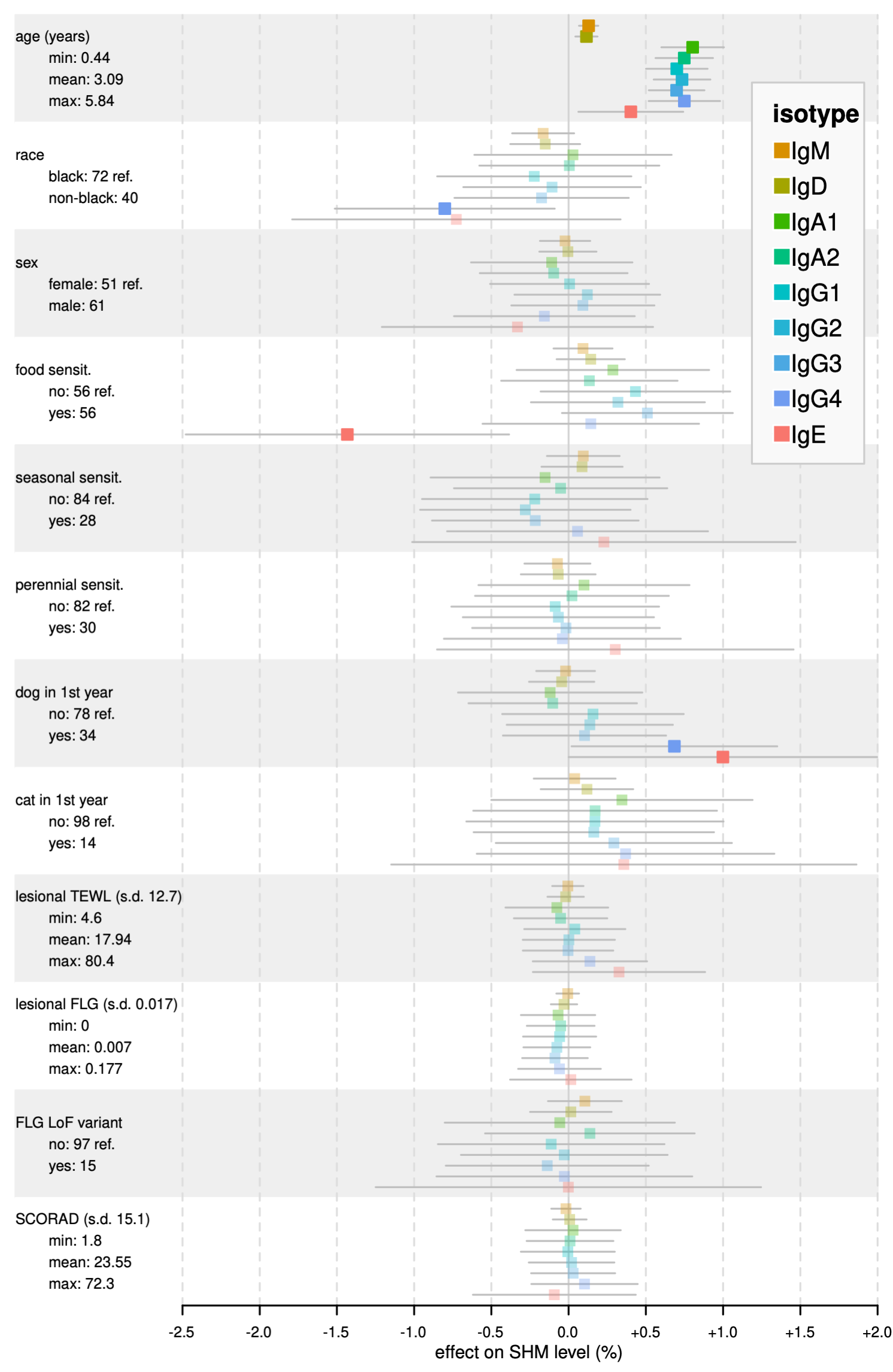

**Fig. S2:** **Linear modeling of SHM, skin barrier function, AD severity and genetic risk.** A sub-cohort (*n=112*) with various measures of AD severity and skin barrier function was analyzed. No significant differences in SHM were seen based on transepidermal water loss (TEWL), expression of filaggrin (FLG), presence of loss-of-function variants in FLG, or SCORing Atopic Dermatitis (SCORAD) measurements. TEWL was measured at lesional sites and values were Z-score normalized before incorporating into the model. FLG expression was measured in keratinocytes and expression was Z-score normalized before incorporating into the model. SCORAD results were Z-score normalized before incorporating into the model.

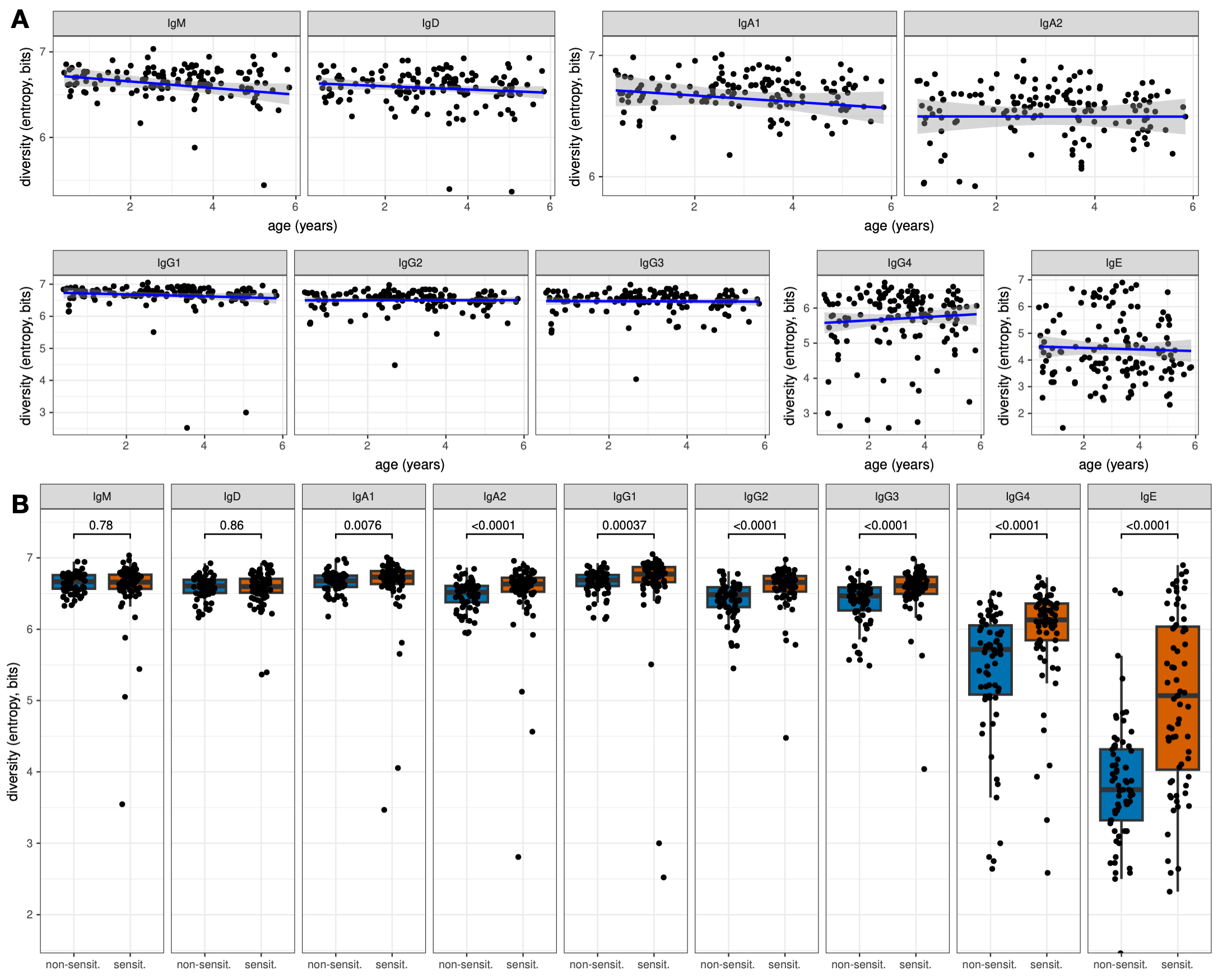

**Fig. S3: Repertoire diversity across all isotype compartments.** **(A)** No statistically significant correlation with age and repertoire diversity was observed for any isotype, although IgM was approaching significance (*p*-value= 0.055 for IgM, *p*-value> 0.11 for others). **(B)** The IgE compartment, the isotype directly involved in allergy and atopic disease, showed a marked increase in diversity in food allergen sensitized subjects as compared to subjects with no food sensitizations. The IgE diversity of food allergen sensitized subjects begin to approach the diversity observed in the other isotypes.

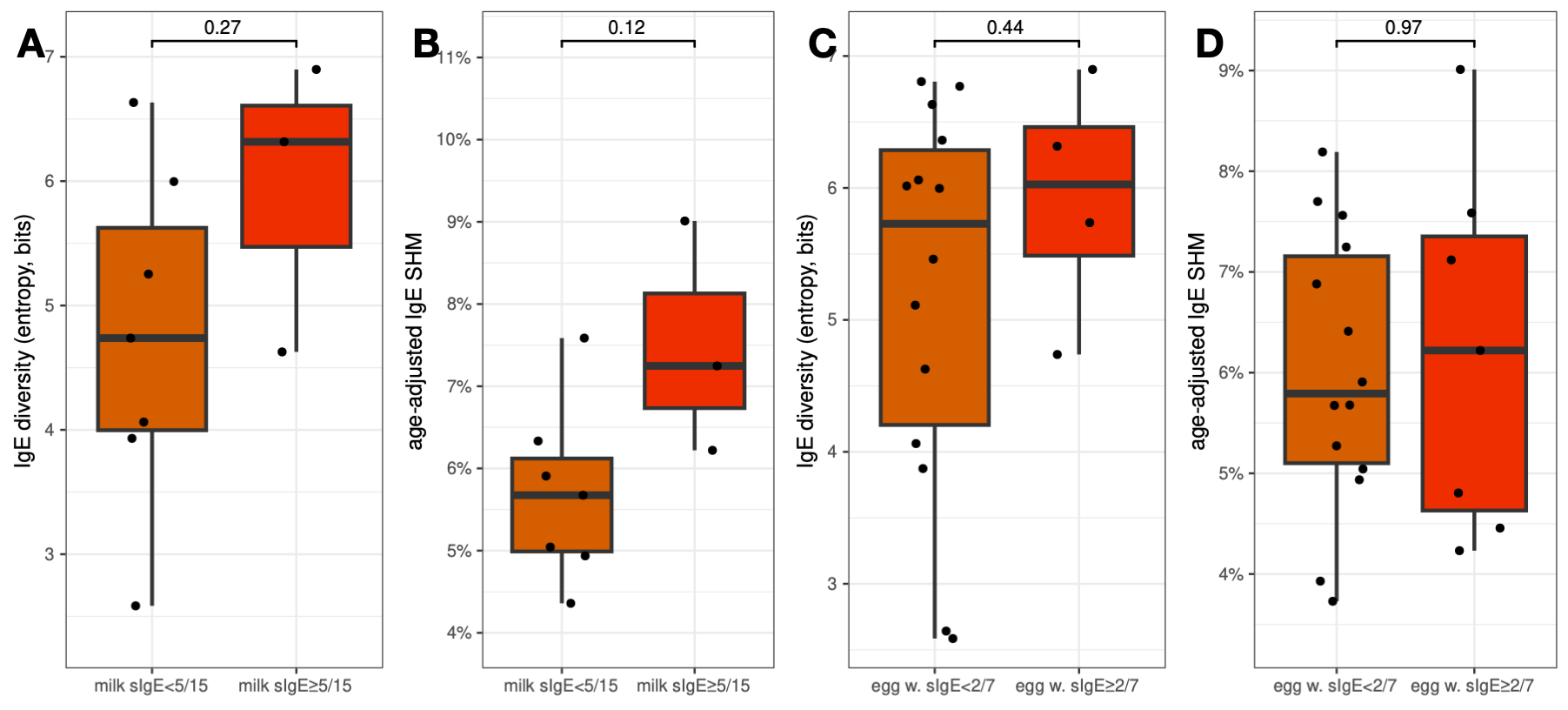

**Fig. S4: IgE diversity and age-adjusted IgE SHM is not significantly altered in subjects highly likely to be milk or egg white allergic.** IgE SHM levels were age-adjusted to the levels they would be expected to have at 3 years of age (the middle of the age range of this cohort) using a linear model (Fig. 1A) **(A)** Subjects with milk sIgE levels above the age-specific 95% positive predictive threshold had age-adjusted IgE SHM levels that trended higher but, with three such subjects, no conclusions can be drawn^16,18^. **(B)** Likewise, age-adjusted IgE SHM trended higher in those likely to be milk allergic. **(C)** Subjects with egg white sIgE levels above the age-specific 95% positive predictive threshold exhibited no statistically significant differences in IgE diversity or **(D)** age-adjusted IgE SHM^16,19^ but with such few subjects, no strong conclusions can be drawn.

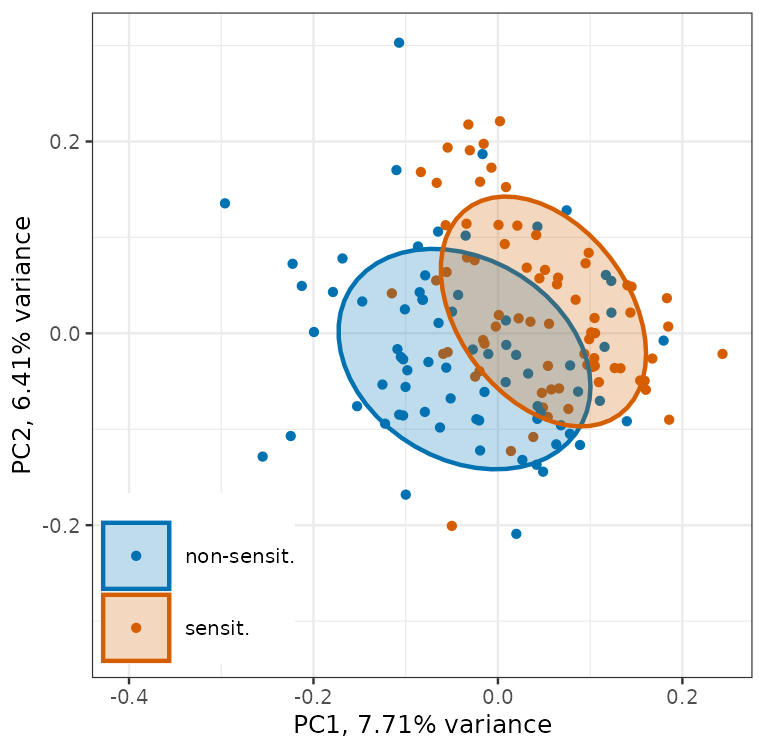

**Fig. S5: Component analysis (PCA) of V-segment usage.** PCA was used on per-subject V-segment clonal usage frequencies in the 5 isotypes of each subject. **(A)** Scatter plot showing the two dimensions which capture the largest trends in the data, PC1 and PC2. Points are colored by food allergen sensitization status and the ellipses show the confidence level of one standard deviation of a multivariate t-distribution around the data. Although PCA does not take phenotype into account, the axes of highest variance (PC1 and PC2) separated food allergen sensitized subjects from subjects with no food allergen sensitizations.
